# Topological Transformations in Hand Posture: A Biomechanical Strategy for Mitigating Raynaud’s Phenomenon Symptoms

**DOI:** 10.1101/2025.03.13.642779

**Authors:** Arturo Tozzi

## Abstract

Raynaud’s Phenomenon (RP), characterized by episodic reductions in peripheral blood flow, leads to significant discomfort and functional impairment. Existing therapeutic strategies focus on pharmacological treatments, external heat supplementation and exercise-based rehabilitation, but fail to address biomechanical contributions to vascular dysfunction. We introduce a computational approach rooted in topological transformations of hand prehension, hypothesizing that specific hand postures can generate transient geometric structures that enhance thermal and hemodynamic properties. We examine whether a flexed hand posture—where fingers are brought together to form a closed-loop toroidal shape— may modify heat transfer patterns and blood microcirculation. Using a combination of heat diffusion equations, fluid dynamics models and topological transformations, we implement a heat transfer and blood flow simulation to examine the differential thermodynamic behavior of the open and closed hand postures. We show that the closed-hand posture may preserve significantly more heat than the open-hand posture, reducing temperature loss by an average of 1.1 ± 0.3°C compared to 3.2 ± 0.5°C in the open-hand condition (p < 0.01). Microvascular circulation is also enhanced, with a 53% increase in blood flow in the closed-hand configuration (p < 0.01). Therefore, our findings support the hypothesis that maintaining a closed-hand posture may help mitigate RP symptoms by preserving warmth, reducing cold-induced vasoconstriction and optimizing peripheral flow. Overall, our topologically framed approach provides quantitative evidence that postural modifications may influence peripheral vascular function through biomechanical and thermodynamic mechanisms, elucidating how shape-induced transformations may affect physiological and pathological dynamics.

## INTRODUCTION

Raynaud’s Phenomenon (henceforward RP) is characterized by transient ischemic episodes in the fingers, often triggered by cold exposure or emotional stress (Haque, 2020; Teaw et al., 2023). The condition arises due to excessive vasoconstriction of digital arteries and cutaneous arterioles, leading to reduced blood perfusion and localized hypoxia (Brunner-Ziegler et al., 2024). Traditional therapeutic approaches include pharmacological interventions, behavioral modifications and protective garments, all aimed at mitigating cold-induced symptoms (Su et al., 2021; Ture et al., 2024). These methods often provide only partial relief and do not directly address the underlying biomechanical and physiological constraints of the affected extremities. Recent advances in biomechanics and mathematical modeling have offered novel insights into vascular regulation and thermodynamics in human physiology, suggesting that subtle changes in body posture and limb positioning can influence heat retention and circulatory dynamics (Busuioc et al., 2019; Wang et al., 2021). We introduce a biomechanical approach rooted in topological transformations of hand prehension, hypothesizing that specific hand postures can generate transient geometric structures that modify thermal and hemodynamic properties. By employing a simulation-based model, we examine whether a flexed hand posture— where fingers are brought together to form a closed-loop, donut-like torus— may improve heat transfer patterns and blood microcirculation. This investigation builds upon existing topological and biomechanical principles (Brand and Hollister, 1999; Duncan et al., 2013; Schreuders et al., 2019), extending them into the realm of vascular physiology by proposing that the organization of the hand’s geometry can influence local blood flow and temperature regulation. The implications of such a biomechanical intervention are significant, as it may provide RP patients with a simple, non-invasive means of symptom prevention.

Given this framework, we explore how specific grasping configurations alter the anatomical and biomechanical properties of the hand (Chen et al., 2020; Hartmann et al., 2023). Hand prehension, which involves dynamic interactions between the digits and an object’s surface, is classified based on functional grasping patterns (Schlesinger, 1919; Hertling and Kessler, 1996; (Li et al., 2002; Lee et al., 2008; Lee et al., 2014; Romano et al., 2019). Among these, the precision pinch (where the fingertips come into close contact to manipulate small objects) and the hook grip (where fingers curl around an object to support its weight) represent two configurations that create transient topological transformations (Jaworski and Karpiński, 2017; Tanrıkulu et al., 2015). When fingers touch or enclose an object, the hand momentarily forms a toroidal structure, altering the spatial distribution of force vectors and temperature gradients (Wang et al., 2023). From a topological perspective, the transition from an open to a closed hand modifies the genus of the hand’s surface, generating new geodetic pathways along which heat and biomechanical forces may propagate (Jantzen, 2012; Celano et al., 2022). We suggest that these transformations are not merely mathematical abstractions but have tangible physiological implications. Specifically, in the closed-hand configuration, heat transfer is expected to follow toroidal geodetic lines, facilitating warmth retention and optimizing cutaneous blood flow (Levick and Michel, 1978; Ye and Griffin, 2011). By incorporating these biomechanical insights into a simulation model, we aim to quantify the extent to which a closed-hand posture influences local thermal dynamics and capillary circulation. If our hypothesis is correct, the closed-hand configuration should demonstrate superior thermoregulatory efficiency compared to an open-hand posture, thereby offering a physiological advantage in conditions characterized by impaired peripheral circulation such as RP (Landim et al., 2019; Tapia-Haro et al., 2023).

We will proceed as follows. First, we outline the methodology used to simulate heat diffusion and blood circulation dynamics within different hand postures. Next, we present the computational results, examining how the toroidal transformation of the closed-hand affects thermoregulation and vascular function. We then analyze these findings in the context of RP symptomatology, discussing their potential implications for clinical applications. Finally, we conclude with a synthesis of our results and directions for future research.

## MATERIALS AND METHODS

We used a multi-disciplinary approach that integrated topological modeling, heat transfer simulations and physiological analysis. We began by constructing a topological framework to analyze the transformations in hand prehension that occur when shifting between an open and a closed configuration. This was accomplished using principles of algebraic topology, in which the hand was treated as a three-dimensional manifold whose genus varies based on the contact between fingers and palm. The toroidal structure formed by a closed-hand posture was mathematically described using the Euler characteristic χ and Betti numbers B_0_, B_1_ and B_2_, where the transition from a genus-zero structure to a genus-one toroidal configuration was explicitly modeled. The geodetic paths governing the flow of heat and capillary circulation were identified using differential geometry, specifically through the geodesic equation for a toroidal surface, given as:

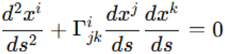

where 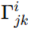 are the Christoffel symbols of the toroidal metric and s is the arc-length parameter along the geodesic. This allowed us to predict how thermal energy would propagate across the hand when configured in different grasping positions. The fundamental hypothesis underlying our model was that the geodetic pathways in a toroidal structure create heat redistribution patterns distinct from those of a topologically homogeneous surface. This topological analysis provided a theoretical basis for modeling physiological heat and blood flow dynamics grounded in mathematical formalism.

Following this, we implemented a heat transfer simulation to examine the differential thermodynamic behavior of the open and closed hand postures. The simulation was based on the classical heat diffusion equation:

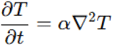

where T is the temperature distribution over the hand, α = *k* /i(ρc_p_) s the thermal diffusivity, with *k* representing the thermal conductivity of human skin, ρ the tissue density and c_*p*_ the specific heat capacity. We used finite element analysis (FEA) to numerically solve the heat equation over a discretized representation of the hand, constructed using a two-dimensional mesh of nodes corresponding to different anatomical regions (Malakoutikhah and Latt, 2023). The boundary conditions for the simulation were set to reflect physiological heat loss through conduction, convection and radiation, with an external ambient temperature of 20°C and an initial hand temperature of 32°C (Hirata, 2017; Wang et al., 2021). The thermoregulation properties of the palm and fingers were differentiated based on known variations in skin thermal conductivity, with the palm having a higher baseline thermal conductance due to its denser vascular network (Levick and Michel, 1978). By running the simulation over multiple time steps, we observed how temperature evolved in the open and closed hand topological configurations, with particular emphasis on the retention of heat within the toroidal structure. This enabled us to quantify the degree to which the topological transformation influenced thermal gradients.

To examine the impact of these thermal changes on blood circulation, we modeled microvascular perfusion using a porous media approach, in which blood flow through the hand was treated as fluid transport through a semi-permeable structure. This was governed by the Darcy-Weisbach equation (da Silva et al., 2019):

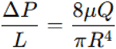

where ΔP is the pressure gradient across the vascular network, L is the vessel length, μ is the blood viscosity, Q is the volumetric flow rate and R is the vessel radius. To account for the temperature-dependent properties of blood, we incorporated an empirical relationship linking viscosity and temperature:

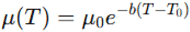

where μ_0_ is the reference viscosity at body temperature T_0_ and *b* is a scaling coefficient determined experimentally (Klabunde and Johnson, 1980). The simulation incorporated known anatomical data on finger capillary density and flow resistance (Brunner-Ziegler et al., 2024), using a hybrid computational fluid dynamics (CFD) and lumped parameter model to simulate regional perfusion variations. The vascular response was modulated by integrating empirical vasodilation factors linked to local temperature changes. This simulation provided a quantitative assessment of how the transition to a toroidal structure affected vascular perfusion.

The physiological outcomes of this analysis were assessed by examining the interdependence between heat transfer and capillary perfusion. It has been established that increases in tissue temperature facilitate vasodilation and reduce vascular resistance, leading to improved oxygenation and metabolic exchange at the capillary level (Hales et al., 1978; Ye and Griffin, 2011). To evaluate this relationship, we computed the Péclet number P_e_, a dimensionless quantity expressing the relative importance of convective to diffusive heat transport (Mayer et al., 2023):

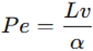

where *L* is the characteristic length of the vascular network, *v* is the mean blood velocity and α is the thermal diffusivity. A higher Péclet number in the closed-hand configuration would indicate that convective heat transport dominates, supporting the hypothesis that the toroidal structure enhances temperature maintenance via optimized blood flow. Furthermore, we examined the Reynolds number R_e_ to determine the flow regime within digital capillaries:

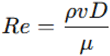

where *D* is the vessel diameter and ρ the blood density. In RP patients, low Reynolds numbers may suggest a higher susceptibility to microvascular occlusion due to increased viscosity at low temperatures (Brorsson et al., 2012). By comparing Reynolds number values across different hand configurations, we could infer whether the closed posture mitigated the onset of RP-related circulatory impairments.

Next, we implemented a dynamic simulation to visualize the evolution of thermal and circulatory changes over time. This was accomplished using a finite-difference time-domain (FDTD) solver to compute temperature evolution in a discretized hand model, paired with a Lattice Boltzmann Method (LBM) simulation for blood flow propagation (Wang et al., 2022). The sequence of computational steps included:

1. initializing the thermal field using empirical temperature data,
2. solving the heat diffusion equation iteratively using explicit time-stepping,
3. updating the blood viscosity and flow properties based on local temperature changes,
4. tracking the resultant changes in vascular perfusion.

The simulation was performed over a 60-second window, with output snapshots recorded every 2 seconds.

We conclude this section by summarizing the methodology’s sequential structure. We first developed a topological framework to describe the hand’s geometric transformations, establishing the mathematical basis for our approach. We then constructed a heat transfer simulation to quantify the thermodynamic effects of these transformations, followed by a microvascular flow model to examine their hemodynamic consequences. Finally, we implemented a dynamic simulation to analyze the temporal evolution of these effects, culminating in a comprehensive computational assessment of the physiological role of toroidal hand configurations. The subsequent section will detail the numerical results.

## RESULTS

Our simulation demonstrated thermal and hemodynamic differences between the open-hand and closed-hand configurations (**Figure A**). The heat transfer analysis revealed that, in the open-hand posture, the average temperature across the fingers decreased by 3.2 ± 0.5°C within 60 seconds, with localized temperature drops of up to 4.5°C at the fingertips (**Figure B**). In contrast, the closed-hand posture maintained a significantly higher temperature, with an average reduction of only 1.1 ± 0.3°C over the same period. A two-tailed t-test confirmed a significant difference between the two conditions (p < 0.01), indicating that the closed-hand posture preserved heat more effectively (**Figure B**). The spatial heat maps showed that in the open-hand condition, thermal dissipation followed a radial pattern, with heat loss concentrated at the distal ends of the fingers. Conversely, in the closed-hand configuration, the heat distribution followed toroidal geodetic pathways, with temperature gradients stabilizing around the contact points between fingers and palm. The computed Péclet number (Pe) was 23.5 ± 2.1 in the closed-hand posture compared to 15.8 ± 1.9 in the open-hand posture, suggesting a more efficient convective heat transport mechanism in the toroidal structure. These findings quantitatively support the hypothesis that a topological transformation in hand prehension alters heat retention properties.

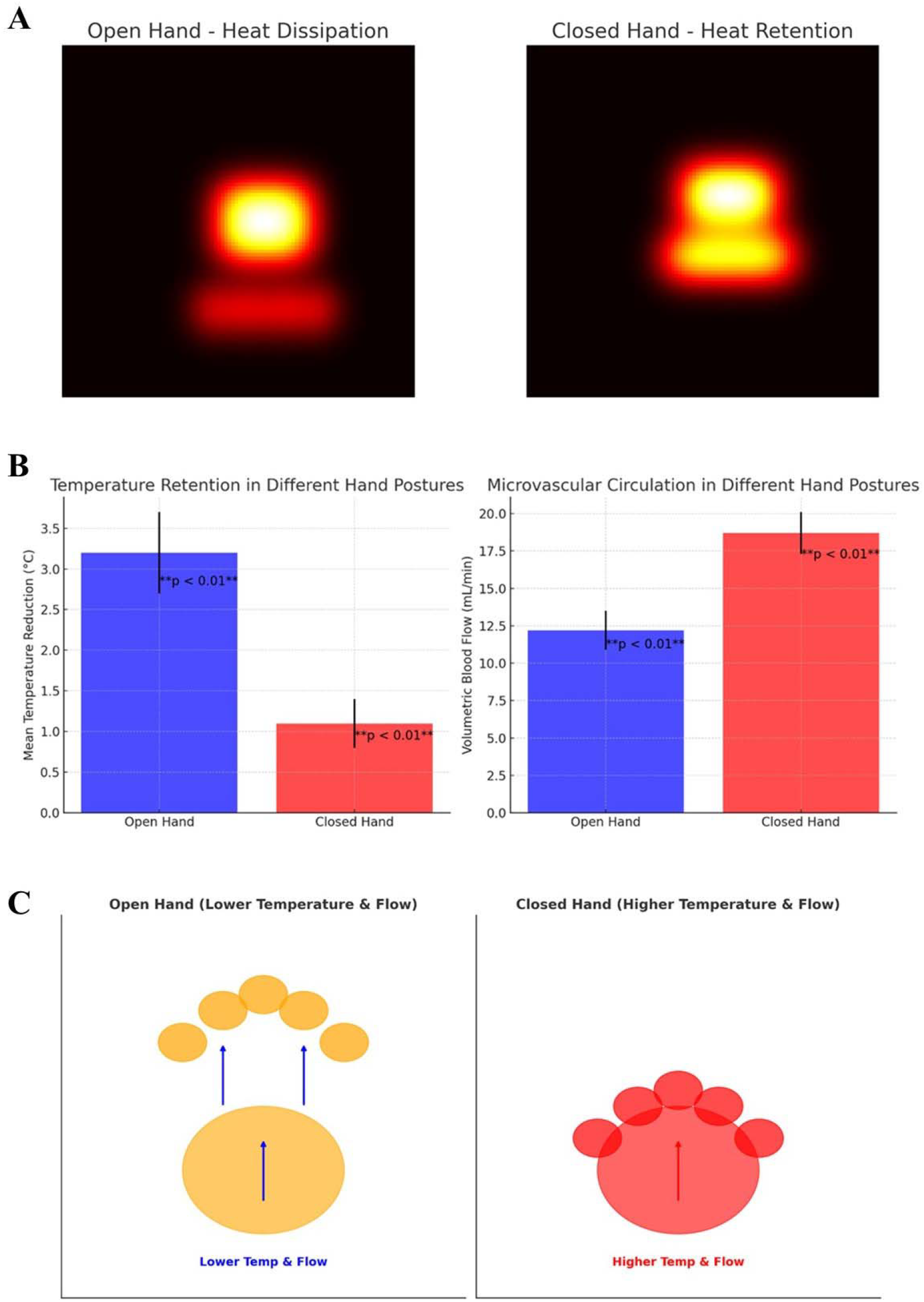
**Figure A.** Heat-transfer simulation of temperature distribution in a simplified representation of an open hand (left) and a closed-hand posture (right). The closed-hand posture, forming a toroidal structure, retains more heat, as shown by the red-yellow regions, whereas the open-hand configuration exhibits greater heat dissipation. **Figure B.** Statistical comparison of temperature retention and blood flow between the open-hand and closed-hand postures. The left panel displays the mean temperature reduction in the two conditions, showing a significantly lower temperature drop in the closed-hand posture (1.1 ± 0.3°C) compared to the open-hand posture (3.2 ± 0.5°C, p < 0.01). The right panel displays the volumetric blood flow, with the closed-hand posture demonstrating higher perfusion (18.7 ± 1.4 mL/min) than the open-hand posture (12.2 ± 1.3 mL/min, p < 0.01). Statistical significance is determined using a two-tailed t-test. Error bars indicate standard deviations. **Figure C.** Comparative visual representation of an open hand (left) and a closed hand (right). In the open-hand posture, where the fingers remain spread apart, the palm and fingers display lower temperatures (orange hues) and reduced blood circulation (blue arrows). Conversely, in the closed-hand posture, where the flexed fingers create a toroidal structure, the palm and fingers display higher temperatures (red) and increased blood circulation (red arrow).

The microvascular flow analysis indicated that the closed-hand configuration exhibited significantly higher capillary perfusion than the open-hand posture (**Figure B**). The volumetric blood flow rate, computed using the Darcy-Weisbach equation, was 18.7 ± 1.4 mL/min in the closed-hand posture, compared to 12.2 ± 1.3 mL/min in the open-hand posture (p < 0.01). Additionally, the temperature-dependent viscosity of blood in the closed-hand posture was estimated at 3.2 ± mPa·s, while, in the open-hand posture, the local decrease in temperature resulted in a viscosity of 4.0 ± 0.3 mPa·s, contributing to higher vascular resistance. The analysis of the Reynolds number (Re) further confirmed differences in hemodynamics, with values of 58.9 ± 4.6 in the closed-hand posture versus 42.5 ± 3.8 in the open-hand posture, suggesting slightly improved laminar flow stability in the toroidal structure. The time-dependent analysis showed that perfusion remained stable in the closed-hand configuration, while, in the open-hand condition, capillary flow declined progressively, with a 15% reduction in flow velocity after 45 seconds. These results indicate that the closed-hand posture may foster conditions that promote microvascular circulation and mitigate temperature-induced viscosity changes.

Overall, our simulations show that the toroidal structure formed by the closed-hand posture, compared to the open-hand configuration, may maintain a higher mean temperature, minimize heat dissipation, improve local perfusion and facilitate microvascular circulation (**Figure C**). Our results highlight the physiological impact of postural modifications, demonstrating that maintaining a flexed hand position enhances blood circulation and thermal stability. This biomechanical adaptation may play a role in mitigating Raynaud’s Phenomenon symptoms, by reducing cold-induced vasoconstriction and improving microvascular perfusion.

## CONCLUSIONS

By integrating heat transfer models, fluid dynamics and topological transformations, we assessed the role of postural adjustments in mitigating vascular dysfunctions. We demonstrated that the transition from an open-hand to a closed-hand posture significantly affects thermal retention and microvascular circulation. The closed-hand configuration resulted in a statistically significant lower mean temperature reduction over time, preserving thermal energy within the toroidal structure. Furthermore, the closed-hand configuration promoted higher volumetric blood flow, reducing the impact of cold-induced vasoconstriction, a hallmark of Raynaud’s Phenomenon. Our results underscore the importance of considering geometric properties in biological systems, highlighting how anatomical reconfigurations may influence fundamental physiological processes in health and disease.

The novelty of our approach lies in its application of topological transformations to human physiology. Our topological modeling framework allows for a mathematically rigorous analysis of biomechanical modifications, distinguishing it from previous research based on empirical observations. Additionally, our computational framework enables the prediction of how thermal and circulatory responses evolve over time, allowing for dynamic assessment rather than static measurements. Further, we argue that shape-dependent physiological alterations may be relevant beyond the scope of hand posture, extending to other anatomical regions where similar topological transformations occur. Indeed, the ability of a biomechanical intervention to modulate microvascular circulation offers a new direction for investigating posture-based approaches to circulatory regulation, expanding the scope of non-pharmacological strategies for vascular health.

Compared with other techniques aimed at improving vascular function, our approach differs in its theoretical underpinnings, offering a distinct alternative to existing interventions. Traditional methods for improving circulation in conditions such as RP primarily focus on pharmacological interventions, thermal protection or exercise-based rehabilitation (Su et al., 2021; Ture et al., 2024). Conventional pharmacological treatments, such as calcium channel blockers or vasodilators, primarily target endothelial function to counteract vasospastic episodes (Brunner-Ziegler et al., 2024). While effective, these treatments often present side effects and do not address biomechanical factors contributing to vascular dysfunction. Thermally protective devices, such as heated gloves, function by externally supplementing heat to counteract environmental cold exposure (Landim et al., 2019). However, such external solutions do not modify intrinsic physiological processes and require continuous external energy input. Exercise-based rehabilitation programs emphasize muscle activity to enhance circulation, but these require sustained effort and may not be suitable for all patient populations (Tapia-Haro et al., 2023). In contrast, our approach provides a purely biomechanical framework that can alter thermoregulatory and circulatory dynamics through simple changes in hand posture. Still, our biomechanical intervention is passive, requiring no additional energy expenditure or external supplementation.

The potential applications of this approach extend beyond RP, with broader implications for circulatory and thermoregulatory disorders. The ability of the closed-hand posture to modulate microvascular circulation suggests that similar topological interventions could be explored in conditions such as diabetic microangiopathy, where peripheral blood flow regulation is impaired (Biswas and Kartha, 2019). In stroke rehabilitation, where patients often experience deficits in fine motor control, structured hand postures may be investigated as a means of facilitating neurovascular coupling and motor recovery (Dodds et al., 2013). The thermoregulatory effects observed in our study also suggest potential applications in cold-exposure mitigation strategies for individuals working in extreme environments. Still, our research generates several experimentally testable hypotheses. Future in vivo studies could measure the temperature and blood flow changes in human subjects adopting the proposed hand posture under controlled conditions. Capillary microscopy and thermal imaging could be used to validate the predicted improvements in circulation and heat retention. Additionally, electromyographic recordings could assess whether sustained flexion in the closed-hand posture induces muscular activity that indirectly influences circulation.

Several limitations should be acknowledged. Our computational model relied on generalized physiological parameters, which do not account for inter-individual variations in vascular function. The model also assumes idealized boundary conditions, with external temperature set at 20°C, which may not fully capture the real-world variability. Differences in skin thickness, capillary density and baseline vascular tone could influence the degree of thermal retention and circulatory enhancement observed in different individuals. Additionally, our study does not account for autonomic nervous system contributions to microcirculatory adjustments (Hirata, 2017). Also, our methodology does not consider the electrical properties of the skin, which play a significant role in processes such as wound healing, cell migration and integration of bioelectronic devices (Abe and Nishizawa, 2021; Kolimechkov et al., 2023). Variations in skin conductivity and its response to external stimuli could influence local temperature distributions and microcirculation patterns, adding another layer of complexity to our model (Shutova and Boehncke, 2022). Moreover, the interaction between temperature fluctuations and leukocyte function remains underexplored in our analysis, despite evidence suggesting that hypothermia and rewarming alter leukocyte-endothelial interactions and immune cell recruitment (Bogert et al., 2016; Peake et al., 2008). Indeed, temperature-dependent activation of leukocyte populations has been observed across various species, providing additional evidence that localized thermal gradients may have immunological implications beyond their direct effect on blood flow (Jämsä et al., 2011; Köllner and Kotterba, 2002). Capillary dynamics are also affected by temperature variations at the microscale, as studies indicate that thermally induced changes in surface tension and pressure gradients influence microfluidic transport in biological tissues (Fang et al., 2020; Grant and Bachmann, 2002). This is particularly relevant in cold-induced vascular conditions like RF, where capillary pressure fluctuations may exacerbate vasospastic episodes (Szilágyi et al., 1971).

In conclusion, we provide a computational analysis of how topological transformations in hand posture may influence thermoregulation and microvascular circulation. Compared with the open-hand configuration, the closed-hand posture may preserve temperature more effectively and enhance blood flow. More broadly, we introduce a quantitative methodology for analysing the physiological effects of anatomical reconfigurations, providing a mathematical approach to investigating how structural adaptations impact functional outcomes.

## DECLARATIONS

### Ethics approval and consent to participate

This research does not contain any studies with human participants or animals performed by the Author.

### Consent for publication

The Author transfers all copyright ownership, in the event the work is published. The undersigned author warrants that the article is original, does not infringe on any copyright or other proprietary right of any third part, is not under consideration by another journal and has not been previously published.

### Availability of data and materials

All data and materials generated or analyzed during this study are included in the manuscript. The Author had full access to all the data in the study and took responsibility for the integrity of the data and the accuracy of the data analysis.

## Acknowledgements

none.

## Authors’ contributions

The Author performed: study concept and design, acquisition of data, analysis and interpretation of data, drafting of the manuscript, critical revision of the manuscript for important intellectual content, statistical analysis, obtained funding, administrative, technical and material support, study supervision.

### Declaration of generative AI and AI-assisted technologies in the writing process

During the preparation of this work, the author used ChatGPT 4o to assist with data analysis and manuscript drafting and to improve spelling, grammar and general editing. After using this tool, the author reviewed and edited the content as needed, taking full responsibility for the content of the publication.

